# Neuro-anatomical and neuro-molecular markers in salience network and their integration in machine learning-based prediction of cognitive dysfunction in mild cognitive impairment

**DOI:** 10.1101/2021.12.22.473775

**Authors:** Ganesh B. Chand, Deepa S. Thakuri, Bhavin Soni

## Abstract

Recent studies indicate disrupted functional mechanisms of salience network regions, especially right anterior insula (RAI), left AI (LAI), and anterior cingulate cortex (ACC), in mild cognitive impairment (MCI). However, the underlying neuro-anatomical and neuro-molecular mechanisms in these regions are not clearly understood yet. It is also unknown whether integration of multi-modal neuro-anatomical and neuro-molecular markers could predict cognitive impairment better in MCI. Herein we quantified neuro-anatomical volumetric markers via structural magnetic resonance imaging (sMRI) and neuro-molecular amyloid markers via positron emission tomography with Pittsburgh compound B (PET PiB) in SN regions of MCI (n = 33) and healthy controls (n = 27). We found that neuro-anatomical markers are significantly reduced while neuro-molecular markers are significantly elevated in SN nodes of MCI compared to healthy controls (p < 0.05). These altered markers in MCI patients were associated with their worse cognitive performance (p < 0.05). Our machine learning-based modeling further suggested that the integration of multi-modal markers predicts cognitive impairment in MCI superiorly compared to using single modality-specific markers. Overall, these findings shed light on the underlying neuro-anatomical volumetric and neuro-molecular amyloid alterations in SN regions and show the significance of multi-modal markers integration approach in better predicting cognitive impairment in MCI.

## 1. Introduction

Alzheimer’s disease (AD) is a neurodegenerative condition currently affecting ~6.2 million older population in the United States and ~55 million individuals worldwide, and it causes substantial emotional, physical, and financial impacts (2021). Investigating AD in its early phase—mild cognitive impairment (MCI)—is crucial for understanding the underlying mechanisms. Normal cognition involves the coordinated activities among brain regions (Chand and Dhamala, 2017; Chand et al., 2018; Deco et al., 2011; Power et al., 2011). The three brain regions—right anterior insula (RAI), left anterior insula (LAI), and anterior cingulate cortex (ACC)—form a network often referred to as salience network (SN) (Chand and Dhamala, 2016b; Lamichhane et al., 2016; Seeley et al., 2007). Our recent studies illustrated that SN nodes are functionally impaired in MCI (Chand et al., 2017b; Chand et al., 2017c), and the underlying neuro-anatomical and neuro-molecular mechanisms are not clearly understood. It is also unknown whether taking multi-modal—neuro-anatomical and neuro-molecular—markers combined or separately in SN regions predicts cognitive impairment better in MCI.

The SN nodes play crucial roles in normal brain mechanisms (Menon, 2015; Uddin, 2015), such as controlling activities in other brain regions (Chand and Dhamala, 2016a; Goulden et al., 2014; Menon, 2011; Sridharan et al., 2008; Wu et al., 2016). AI regions of SN are functionally involved in coordinating activities in many cognitive processes, including reorientation of attention (Ullsperger et al., 2010) and switching between cognitive resources (Uddin and Menon, 2009). ACC is primarily involved in increased cognitive controls and conflict monitoring (Egner, 2009) and regulates neural activities in association with AI regions during difficult tasks (Chand and Dhamala, 2016a). SN regions have been formerly explained in terms of unique cyto-architecture that they contain a special type of neurons named von Economo neurons (VENs) (Allman et al., 2005). Previous functional MRI studies suggest the disrupted functions of SN nodes in MCI (Chand et al., 2017a, b; Chand et al., 2017c). Investigation of the underlying neuro-anatomical and neuro-molecular changes in SN nodes could help identify AD mechanisms specific to those regions and their associated processes. Volumetric measures of structural magnetic resonance imaging (sMRI) are widely used to study neuroanatomical changes (Chand et al., 2020a). Positron emission tomography with Pittsburgh compound B (PET PiB) is one of the commonly used measures to study neuro-molecular changes, especially amyloid beta plaques, a histopathological hallmark of AD (Bilgel et al., 2018; Cohen and Klunk, 2014; Spira et al., 2013). Combined study of neuro-anatomical volumetric and neuro-molecular amyloid markers in SN regions using machine learning methods and whether this multi-modal integration approach predicts cognitive impairment in MCI is not investigated yet.

In this study, we seek to examine the patterns of neuro-anatomical and neuro-molecular markers of SN nodes in MCI and healthy controls. We *hypothesize* that neuro-anatomical and neuro-molecular markers will differ in MCI patients compared to healthy controls. We further *hypothesize* that the changes in these markers will result in poor cognitive performance. We also *hypothesize* that the integration of these multi-modal markers will better predict cognitive impairment compared to single modality-specific markers in MCI.

## 2. Materials and methods

### 2.1. Participants

In this study, data of 33 MCI patients and 27 healthy controls within the age range of 62-91 years were used. These data were assessed from a publicly available OASIS dataset (LaMontagne et al., 2019; Marcus et al., 2010). Informed consent was obtained from each subject according to the local institutional guidelines for human subjects. Anonymized data was shared by the OASIS (https://www.oasis-brains.org/) following the data request procedure. Participants were characterized using the Clinical Dementia Rating (CDR) (Morris, 1993; Morris et al., 2001) as either healthy control status or with ‘very mild’ to ‘mild’ Alzheimer’s disease, referred to as MCI. CDR rates the participants in each of the six domains: memory, orientation, judgment and problem solving, function in community affairs, home and hobbies, and personal care. A global CDR score is derived from individual ratings in each domain based on the collateral source and participant interview. A global CDR of 0 indicates no dementia and CDR of 0.5, 1, 2, and 3 indicates very mild, mild, moderate, and severe dementia, respectively. Healthy controls had CDR of 0 while MCI participants had CDR of either 0.5 or 1. Participants with a primary cause of dementia other than Alzheimer’s disease (e.g., vascular dementia, primary progressive aphasia), active neurologic or psychiatric illness, serious head injury, history of clinically meaningful stroke, and use of psychoactive drugs were excluded. In this study, the Mini-Mental State Examination (MMSE), which measures general cognitive status with scores ranging from 0 (severe impairment) to 30 (no impairment), was used to study neuro-anatomical and neuro-molecular signatures in SN. In this sample, mean age (standard deviation) was 77.30 years (6.68) in MCI and 73.13 years (7.48) in healthy controls. Proportion of female participants was 45.45% in MCI and 55.56% in healthy controls. Mean MMSE (standard deviation) was 26.70 (2.64) in the MCI group and 28.70 (1.64) in the healthy control group.

### 2.2. Data acquisition and processing

Dynamic ^11^C-labeled PET 3D scans were acquired for over 60 minutes (24×5 second frames, 9×20 second frames, 10×1 minute frames, 9×5 minute frames) following an intravenous bolus injection of ^11^C-PiB. Participants were scanned in either ECAT HRplus 962 PET scanner or Biograph 40 PET/CT scanner. T1-weighted brain sMRI was acquired using Siemens 3T TrioTim or 3T Biograph mMR or 1.5T Vision scanner. In TrioTim, acquisition parameters were: echo time (TE) = 0.003 s, repetition time (TR) = 2.4 s, inversion time (TI) = 1 s, flip angle = 8 degrees, slice thickness = 1 mm, and resolution = 256 × 256. In Biograph mMR, acquisition parameters were: TE = 0.00295 s, TR = 2.3 s, TI = 0.9 s, flip angle = 9 degrees, and resolution = 256 × 256. In Vision, acquisition parameters were: TE = 4 ms, TR = 9.7 ms, and flip angle = 10 degrees. **MU**lti-atlas region **S**egmentation utilizing **E**nsembles of registration algorithms and parameters and locally optimal atlas selection (MUSE) (Doshi et al., 2016) was used for segmentation of each individual’s T1-weighted MR images into anatomical regions of interest and for computation of gray matter volumes. By design, MUSE utilizes an ensemble of atlases coming from different scanners, field strengths, and acquisition protocols, and thus this method provides results quite robust to such confounds (Srinivasan et al., 2020). The anatomical label image was transformed from MRI to PET space and PET PiB outcomes were quantified as distribution volume ratios (DVRs) using a simplified reference tissue model with cerebellar gray matter reference region (Bilgel et al., 2018; Zhou et al., 2007). DVRs were accounted for the partial volume effects using parallel level set (PLS) method (Zhu et al., 2021). ROI volumes and DVRs were corrected for the covariates—scanner differences, age, and sex—using our previous harmonization approach (Chand et al., 2020a). In this approach, the coefficients that account for site (scanner), age, and sex effects were computed using data from all participants and applied to each participant to achieve harmonized data.

### 2.3. Machine learning-based prediction of MMSE using SN features

To evaluate whether combined modeling of neuro-anatomical (volumes) and neuro-molecular amyloid markers (DVRs) in SN nodes better predict cognitive impairment of MCI patients, these markers were used as an input features in the kernel-based support vector machines (SVM) (Cortes and Vapnik, 1995) and MMSE scores were predicted. SVM regression modeling was performed using the libSVM package (Chang and Lin, 2011) using 10-folds cross-validation. SVM cross-validation was consistent with our previous study (Chand et al., 2020b). In this approach, participants were divided into 10-folds, SVM model was trained in 9-folds data, and then the trained model was tested on the remaining 1-fold (test) data. This procedure was repeated for all combinations of 10-folds data. Gaussian kernel function was used and SVM model parameters were optimized within each training set using 10-folds nested cross-validation grid search method (Chand et al., 2020b). SVM regression modeling was also performed using SN volumes or SN DVRs alone as input features. SVM regression performance on MMSE prediction was compared among 1) combined DVRs and volumes, 2) only DVRs, and 3) only volumes of SN regions as input features in machine learning modeling.

### 2.4. Statistical analysis

To compare MCI and healthy control groups, a two-sided Wilcoxon rank sum test was used. Spearman correlations were used to examine associations of gray matter volumes vs. MMSE, and DVR vs. MMSE. Spearman correlations were also used to study the relationship between actual MMSE and predicted MMSE. P-values were corrected for multiple comparisons using false-discovery rate (*FDR*) and were considered statistically significant if *FDR-p < 0.05*.

## 3. Results

### 3.1. Comparison of neuro-anatomical and neuro-molecular markers between MCI and healthy controls

The neuroanatomy of the key nodes of SN—RAI, LAI, and ACC—is shown in **Figure 1**.

**Figure 1:**
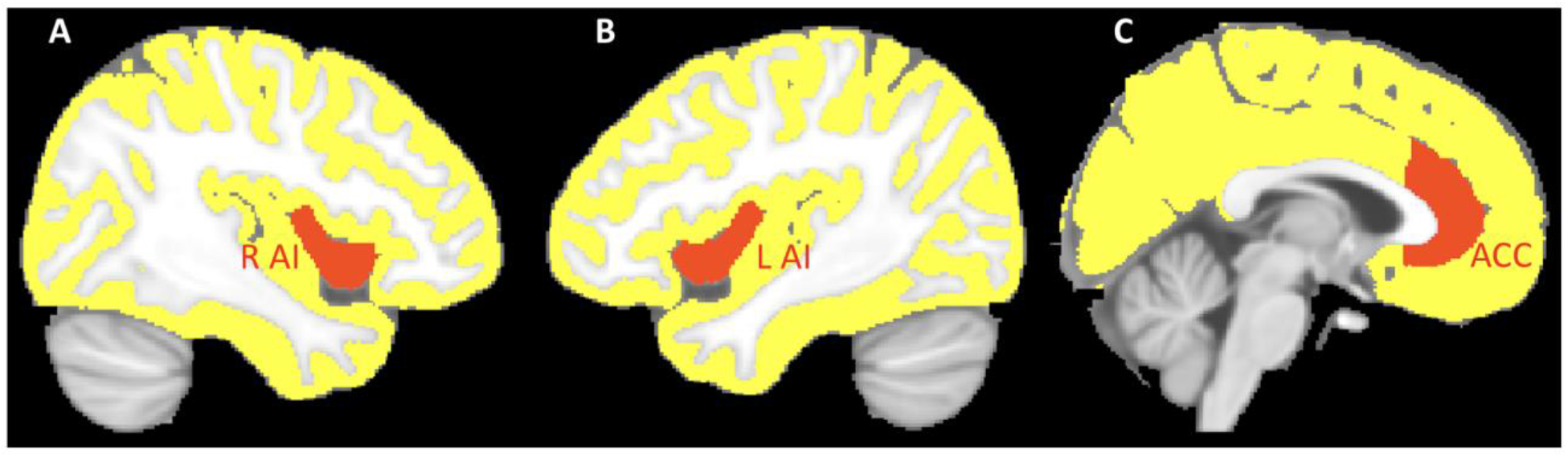
Representation of the key brain regions of salience network (SN): right anterior insula (RAI), left anterior insula (LAI), and anterior cingulate cortex (ACC).

PET PiB outcomes were quantified as DVR in SN nodes and compared between MCI and healthy control groups (**Figure 2**). DVR was significantly elevated in RAI (*FDR-p* < 0.05), LAI (*FDR-p* < 0.05), and ACC (*FDR-p* < 0.05) regions of MCI compared to healthy controls.

**Figure 2:**
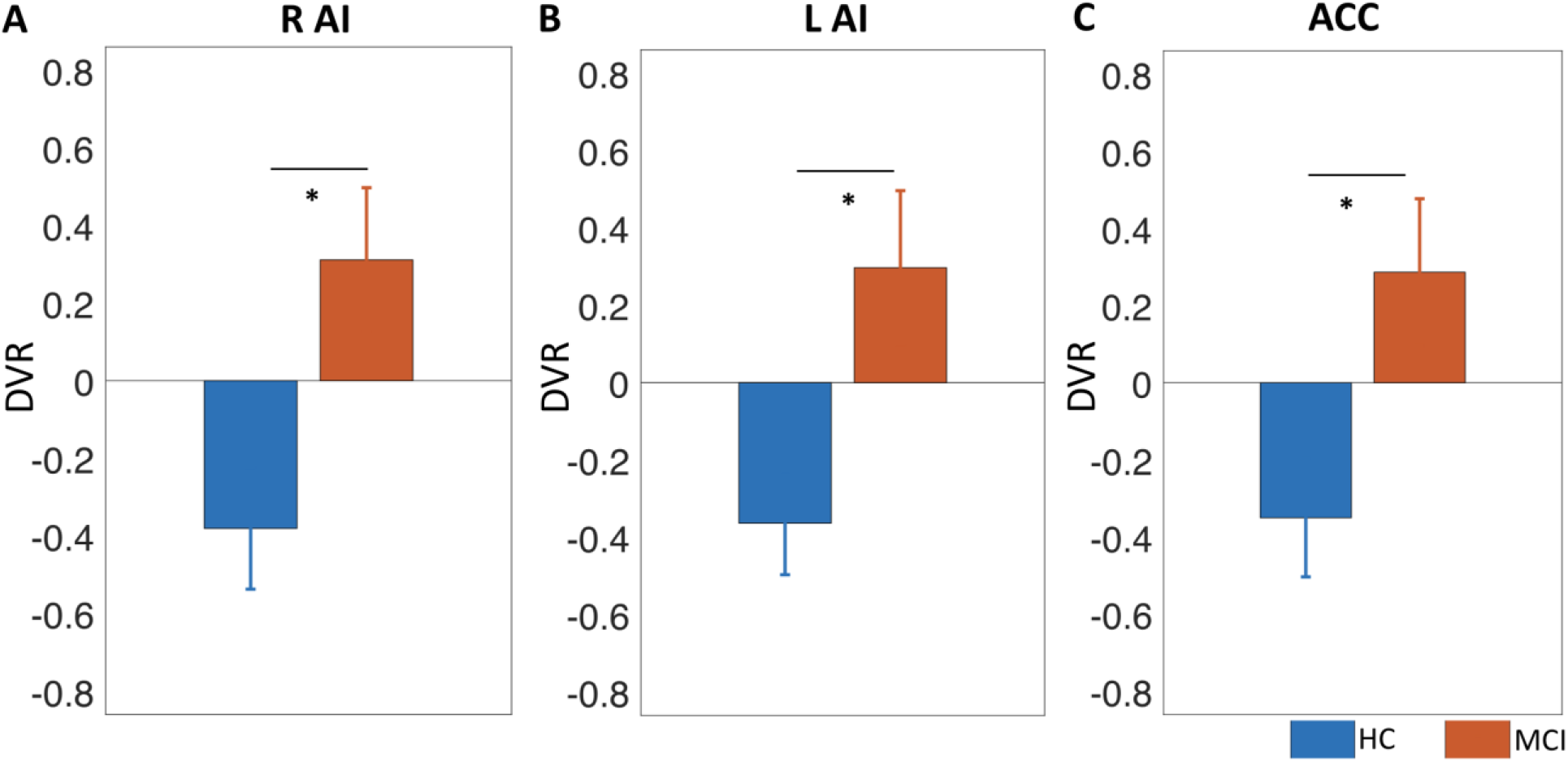
Distribution volume ratio (DVR; z-score) outcome of PET PiB in SN regions: **A-C)** DVR significantly elevated in RAI, LAI, and ACC of MCI compared to healthy controls (HC) (*: p < 0.05; FDR-corrected). Error bars represent standard error of mean (SEM) over participants.

Gray matter volume of each SN node was computed using the MUSE method and compared between MCI and healthy controls (**Figure 3**). Gray matter volume was significantly reduced in RAI (*FDR-p* < 0.05), LAI (*FDR-p* < 0.05), and ACC (*FDR-p* < 0.05) of MCI compared to healthy controls.

**Figure 3:**
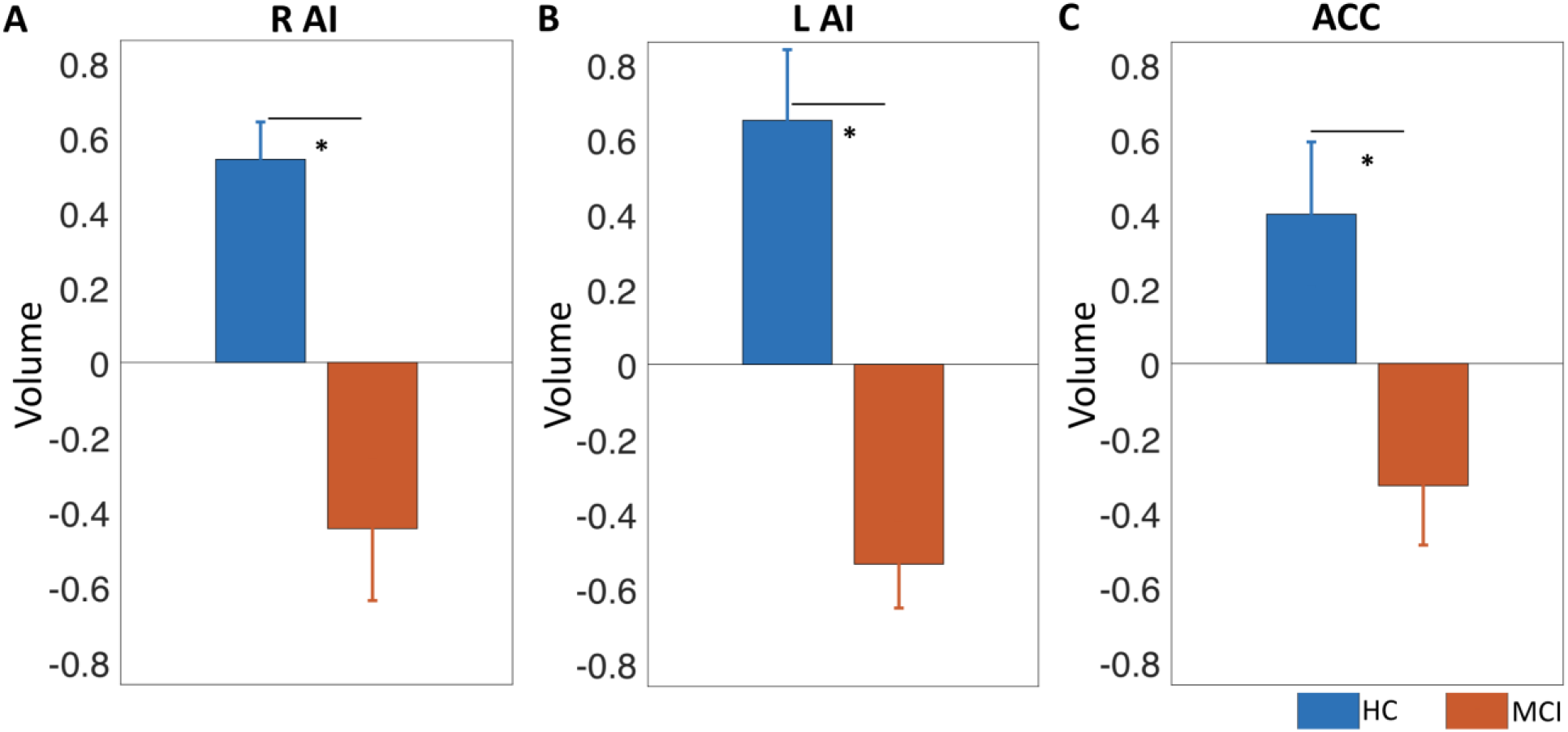
Neuroanatomical volume (z-score) of SN regions: **A-C)** Gray matter volume decreased in RAI, LAI and ACC regions of MCI compared to healthy controls (HC) (*: p < 0.05; FDR-corrected). Error bars represent standard error of mean (SEM) over participants.

### 3.2. Association of neuro-anatomical and neuro-molecular markers with MMSE in MCI

Since MCI patients exhibited a range of MMSE scores, the relationship between DVR of SN nodes and MMSE was studied in MCI (**Figure 4**). MMSE and DVR were negatively associated in RAI (*ρ = −0.51; FDR-p < 0.05*), LAI (*ρ = −0.51; FDR-p < 0.05*), and ACC (*ρ = −0.57; FDR-p < 0.05*) indicating that MCI individuals with elevated DVR has lower MMSE performance.

**Figure 4:**
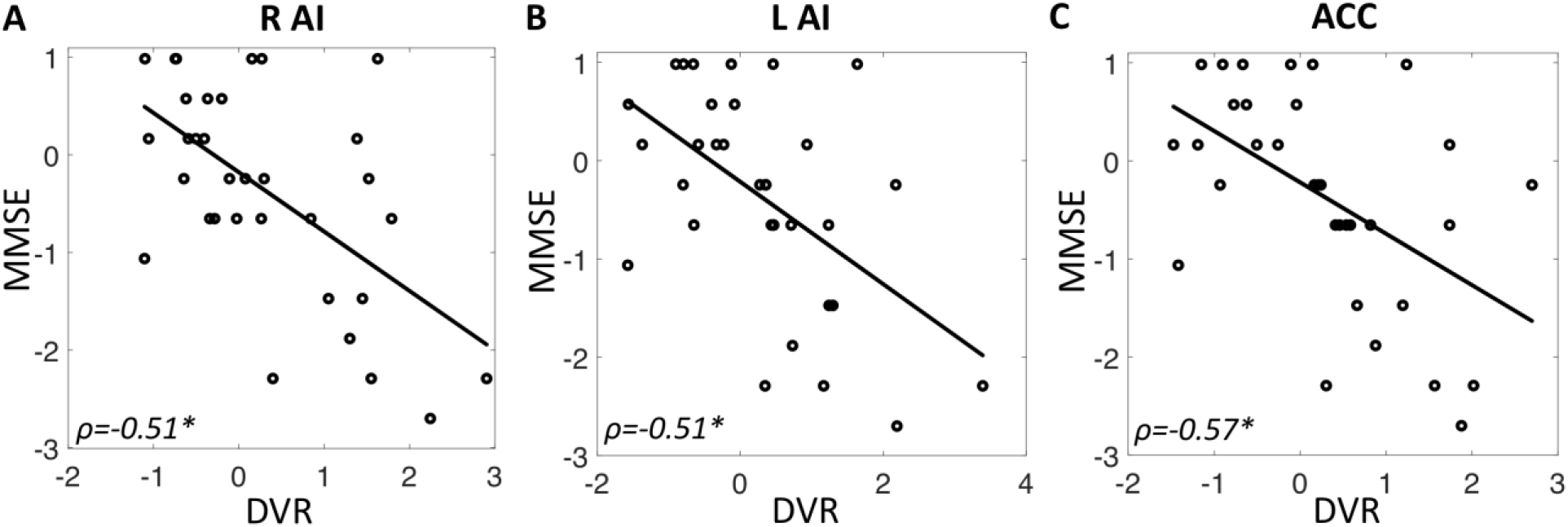
Correlation between DVR (z-score) of SN regions and MMSE (z-score) in MCI: **A-C)** DVR of RAI, LAI and ACC regions of SN negatively correlated with MMSE scores, implying that MCI individuals with higher DVR have higher cognitive impairment (lower MMSE scores) (*: p < 0.05; FDR-corrected).

Correlation between gray matter volume of each SN node and MMSE was also examined in MCI (**Figure 5**). MMSE and gray matter volume was positively correlated in RAI (*ρ = 0.36; FDR-p < 0.05*), but did not reach statistical significance in LAI (*ρ = 0.26; p = 0.15*) and ACC (*ρ = 0.21; p = 0.24*).

**Figure 5:**
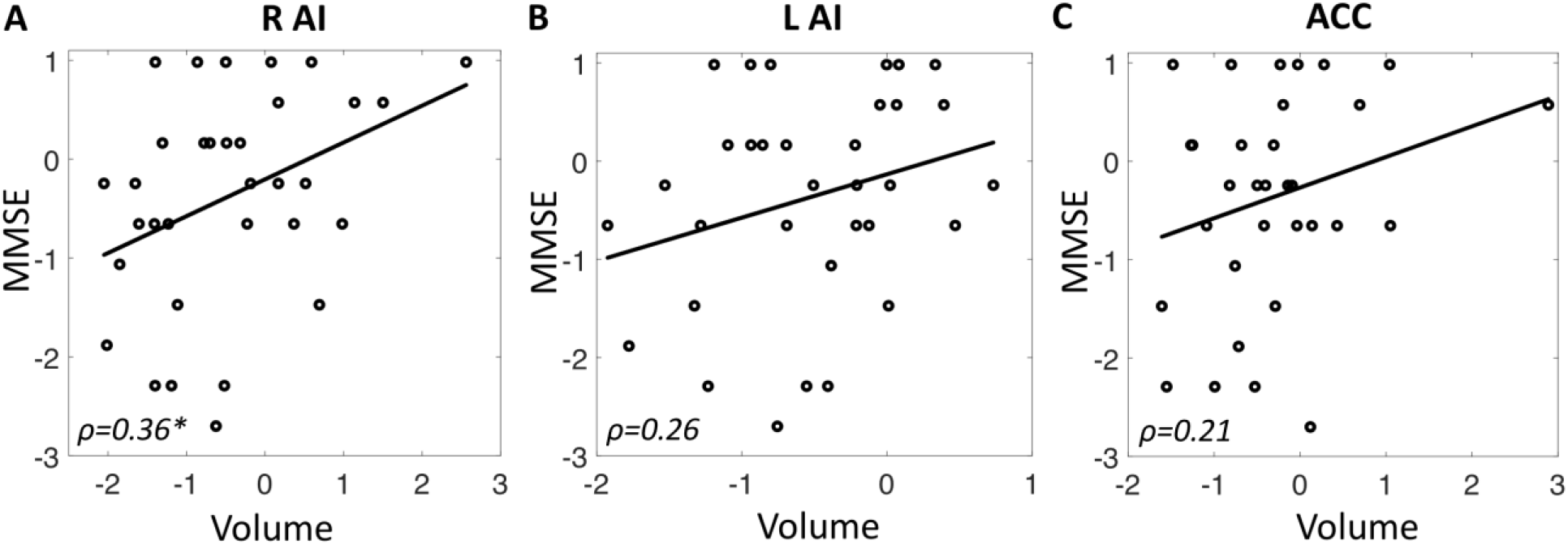
Correlation between neuroanatomical volume (z-score) of SN nodes and MMSE (z-score) in MCI: **A-C)** Gray matter volume of SN regions positively correlated with MMSE scores, implying that MCI patients with lower gray matter volume have higher cognitive impairment (lower MMSE scores) (*: p < 0.05; FDR-corrected).

### 3.3. Machine learning-based prediction of MMSE using neuro-anatomical and/or neuro-molecular markers in MCI

SVM was performed to evaluate the hypothesis that whether an integrated modeling of neuro-anatomical and neuro-molecular signatures superiorly predict MMSE in MCI compared to modeling of individual signatures separately. Using DVR and gray matter volume of SN nodes as input features in SVM, predicted MMSE and actual MMSE were highly correlated in MCI (*ρ = 0.61; FDR-p < 0.05*) (**Figure 6**). Using DVR of SN nodes alone as input features, predicted MMSE and actual MMSE were less correlated in MCI (*ρ = 0.46; FDR-p < 0.05*). Likewise, using only gray matter volume of SN nodes as input features, predicted MMSE and actual MMSE were weakly correlated and were not statistically significant (*ρ = 0.21; p = 0.55*).

**Figure 6:**
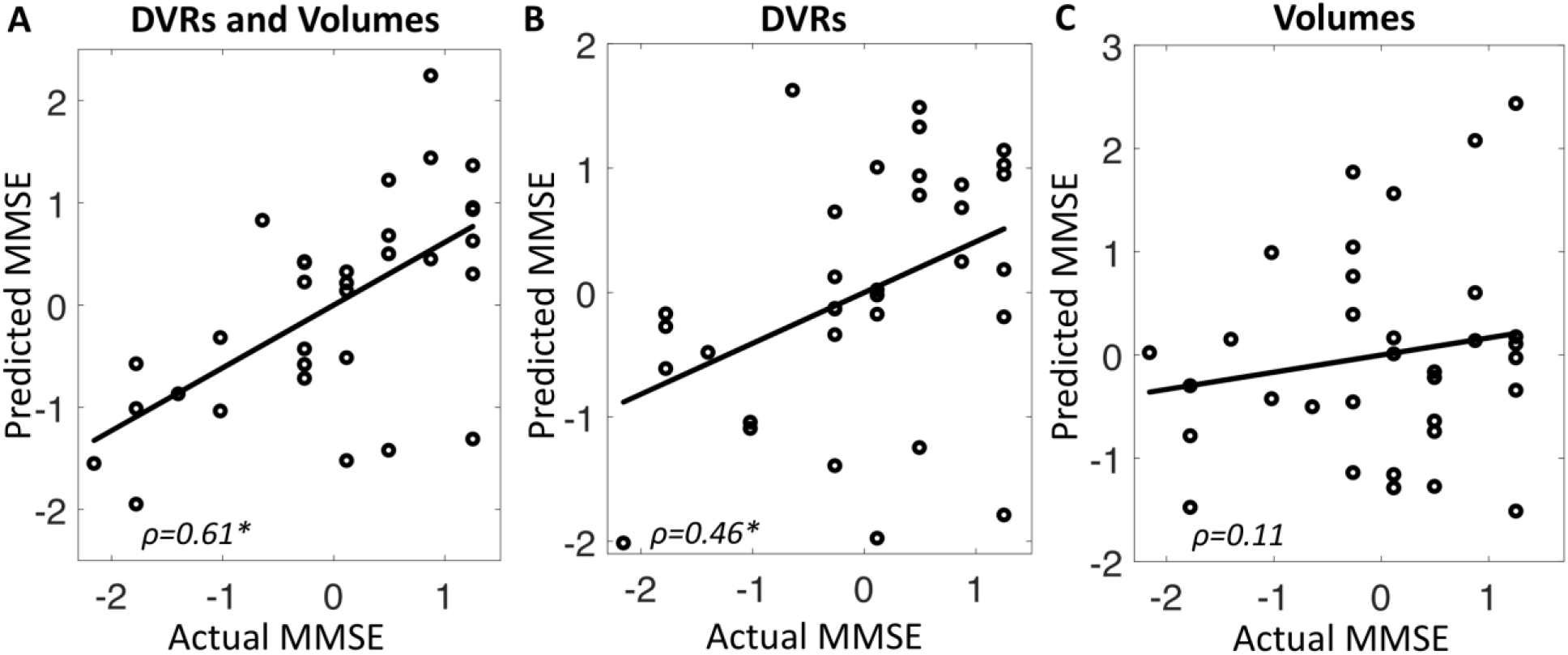
Correlation between actual MMSE (z-score) and predicted MMSE (z-score) in MCI patients using: **A**) DVR and volume markers, **B**) only DVR markers, and **C**) only volume markers as input features in SVM.

## 4. Discussion

This study evaluated multi-modal—neuro-anatomical and neuro-molecular—markers in SN nodes of MCI compared to healthy controls, and assessed their relationships with cognitive impairment. DVR neuro-molecular signatures were significantly elevated while gray matter volume neuro-anatomical signatures were significantly reduced in MCI compared to healthy controls. In MCI patients, higher DVR of SN nodes was associated with lower MMSE (i.e., higher cognitive impairment) while higher gray matter volume was associated with higher MMSE (lower cognitive impairment). Our machine learning approach further demonstrated that the combined modeling of neuro-anatomical and neuro-molecular signatures in SN nodes predict cognitive impairment superiorly (i.e., higher correlation strength between predicted MMSE and actual MMSE) compared to modeling of individual modality-specific signatures.

SN nodes consist of VENs that are relatively larger in size than other neurons such as pyramidal neurons and thus VENs facilitate the coordinated activities among brain regions (Allman et al., 2005; Sridharan et al., 2008). AI connects functionally to other brain regions, including central-executive regions (Vincent et al., 2008) and has direct white-matter connections to ACC (Bonnelle et al., 2012; Jilka et al., 2014), inferior parietal lobe (Uddin et al., 2010), and temporo-parietal junction (Kucyi et al., 2012). These rich structural and functional settings might support AI regions to coordinate activities, such as reorientation of attention (Ullsperger et al., 2010) and switching between cognitive resources (Uddin and Menon, 2009). Similarly, ACC is involved in cognitive controls, conflict monitoring (Egner, 2009), and difficult decision-making tasks (Chand and Dhamala, 2016a). Thus, AI and ACC regions have been argued to be central for active and passive cognitive states in healthy individuals, whereas neuro-anatomical and neuro-molecular impairment to these regions might be associated with the underlying changes to highly sensitive VENs (Allman et al., 2010; Allman et al., 2005; Bonnelle et al., 2012; Butti et al., 2013; Sridharan et al., 2008; Watson et al., 2006). Previous investigations consistently demonstrated altered SN functions in brain diseases (Menon, 2015; Uddin, 2015), including MCI (Chand et al., 2017b; Chand et al., 2017c). Our findings extended prior studies about functional mechanisms of SN (Chand and Dhamala, 2016a; Chand et al., 2017a, b; Goulden et al., 2014; Menon, 2011; Sridharan et al., 2008; Wu et al., 2016) by uncovering both neuro-anatomical and neuro-molecular alterations of SN nodes in MCI. Our results are broadly in line with past studies that report neuro-molecular changes in Alzheimer’s pathology (He et al., 2015; Kim et al., 2021). Our machine learning approach suggested that integration of neuro-anatomical and neuro-molecular multi-modal markers of SN nodes predicts cognitive impairment in MCI superiorly as compared to using modality-specific markers separately.

Although we leveraged multi-modal (neuro-anatomical and neuro-molecular) markers of SN nodes and their integration in machine learning-based prediction of cognitive impairment, it’s worth noting the following limitations. First, the sample size is not large and replication of these findings in larger samples is suggested. Second, we showed a proof-of-concept multi-modal markers integration approach only for SN regions and future studies will need to investigate similar approaches considering other brain regions.

In conclusion, we assessed neuro-anatomical volumetric and neuro-molecular amyloid markers in SN nodes of MCI and healthy controls. We found that volumetric markers are reduced while amyloid markers are elevated in SN nodes of MCI patients. The impairment to these markers was associated with worse cognitive performance of the MCI population. Machine learning modeling suggested that integration of multi-modal markers in SN nodes predict the impairment in MCI population superiorly compared to using single modality-specific markers. Overall, these findings help us better understand the underlying neuro-anatomical and neuro-molecular changes in SN nodes and their potential contributions in predicting cognitive impairment of MCI and/or Alzheimer’s disease population.

## Acknowledgments

Authors acknowledge the publicly available OASIS-3 dataset. Data were provided by OASIS-3 (LaMontagne et al., 2019): Principal Investigators: T. Benzinger, D. Marcus, J. Morris; NIH P50 AG00561, P30 NS09857781, P01 AG026276, P01 AG003991, R01 AG043434, UL1 TR000448, R01 EB009352. Authors also acknowledge the DVR calculation and partial volume correction pipeline (Zhu et al., 2021).

